# CSF1R-Mediated Microglial Engagement Is Required for Stress-Induced MMP-2/9 Activity

**DOI:** 10.64898/2026.06.08.730923

**Authors:** Juan Pablo Taborda Bejarano, Juliana Patricia Tovar, Malika Allen, Jayashree Natarajan, Michael Meyerink, Constanza Garcia-Keller

## Abstract

Stress is a major risk factor for numerous neuropsychiatric disorders and induces enduring synaptic plasticity within the nucleus accumbens core (NAcore), a key brain region involved in reward and stress-related behaviors. Previous studies from our laboratory demonstrated that stress-induced plasticity depends on matrix metalloproteinase (MMP)-2/9-mediated extracellular matrix (ECM) remodeling; however, the upstream cellular mechanisms regulating MMP activation remain unclear. Because microglia regulate neuroimmune signaling, ECM dynamics, and synaptic plasticity, we tested the hypothesis that microglial colony-stimulating factor 1 receptor (CSF1R) signaling contributes to stress-induced MMP-2/9 activation within the NAcore. Male rats received the CSF1R inhibitor PLX3397 prior to acute restraint stress. In vivo fluorescent zymography, immunohistochemistry, and quantitative PCR were used to assess MMP activity, microglial signaling, and inflammatory gene expression. Acute stress increased MMP-2/9 activity enhanced microglial CD68-associated phagocytic signaling, and elevated expression of Csf1r, Tnfa, Cnr2, and Mmp16 within the NAcore. Importantly, CSF1R inhibition attenuated stress-induced increases in MMP-2/9 activity and CD68 immunoreactivity. Combined, these findings identify microglial CSF1R signaling as an upstream regulator of stress-induced ECM remodeling within the NAcore and provide mechanistic insight into how acute stress recruits neuroimmune pathways to remodel reward circuitry.

## Introduction

Stress is a major precipitating factor for numerous neuropsychiatric disorders, including anxiety disorders, post-traumatic stress disorder (PTSD), and substance use disorders[1]. Exposure to acute and chronic stress produces persistent neuroadaptations within reward-related brain regions, particularly the nucleus accumbens core (NAcore)[2-7], a critical structure involved in motivation, reward processing, and stress-related behavioral responses. Previous studies from our laboratory demonstrated that acute restraint stress induces long-lasting cocaine-like structural and functional synaptic plasticity within the NAcore[2, 5, 8]. Similarly, cocaine-induced synaptic plasticity depends on activation of matrix metalloproteinases (MMP)-2 and -9 and subsequent extracellular matrix (ECM) remodeling[9-12]. Because stress and psychostimulants induce overlapping forms of NAcore plasticity, these findings suggest that stress-induced neuroadaptations may also depend on MMP-mediated ECM remodeling. However, the cellular mechanisms that initiate stress-induced MMP activation remain poorly understood.

Microglia, the resident immune cells of the central nervous system, are increasingly recognized as active regulators of synaptic plasticity and circuit remodeling rather than solely immune surveillance cells[13]. In addition to their classical immune functions, microglia regulate ECM composition, cytokine signaling, synaptic organization, and experience-dependent structural plasticity[14-19]. Stress rapidly alters microglial function and promotes neuroimmune signaling within reward-related circuits, suggesting that microglia may contribute to stress-induced synaptic remodeling[20]. One signaling pathway particularly relevant to microglial activation is the colony-stimulating factor 1 receptor (CSF1R), a receptor tyrosine kinase that regulates microglial survival, proliferation, phagocytic activity, and inflammatory signaling[21, 22]. Pharmacological inhibition of CSF1R signaling using PLX3397 (Pexidartinib) has emerged as a powerful strategy for investigating microglial contributions to neurobiological processes[23, 24].

Microglia are also positioned to influence ECM remodeling through the release of cytokines and signaling molecules that regulate MMP activity[18]. Among these, tumor necrosis factor-α (TNFα) has been implicated in stress-induced neuroimmune responses and can promote ECM remodeling and synaptic plasticity[15-17, 24] [19]. In addition, membrane-associated metalloproteinases such as MMP16 can regulate ECM turnover and activation of downstream MMP signaling pathways. Thus, microglial activation represents a plausible upstream mechanism linking stress exposure to MMP-dependent remodeling of reward circuitry.

Here, we tested the hypothesis that microglial CSF1R signaling contributes to stress-induced MMP-2/9 activation and neuroimmune responses within the NAcore. Using in vivo fluorescent zymography, immunohistochemistry, and quantitative PCR, we examined the effects of acute restraint stress on MMP activity, microglial CD68-associated lysosomal phagocytic signaling, and inflammatory gene expression. Furthermore, we determined whether inhibition of CSF1R signaling with PLX3397 attenuates these stress-induced neuroimmune and ECM adaptations. Our findings identify microglial CSF1R signaling as a critical regulator of stress-induced MMP activation and provide evidence that microglia function upstream of ECM remodeling within reward circuitry.

## Methods

### Animal housing

Male Long-Evans rats (250-300g) were double- or triple-housed under a reverse 12:12 hr dark/light cycle. The animals were approximately 8 weeks old (+/-2 weeks). All experiments were done during the dark cycle. Rats received food and water ad libitum. All housing and experimental procedures were conducted in accordance with the Medical College of Wisconsin Institutional Animal Care and Use Committee (IACUC) regulations.

### Acute restraint stress

Male rats were randomly assigned to either the sham or stress groups. The stressed rats were confined in an acrylic flat-bottom restrainer (PLAS Labs, Thomas Scientific, Swedesboro, NJ, USA) for 30 minutes between 8:00 and 13:00 h, then placed in a cage identical to their home cage. The sham animals remained in a similar cage for 30 minutes without restraint. Both groups were euthanized and perfused after this period[2, 5].

### In-vivo zymography analysis and microinjection

MMP activity is measured using an in vivo zymography assay as described previously [10, 12]. Rats were stereotaxically implanted with a unilateral cannula aimed above the NAcore (AP +1.6, ML ±1.6, DV −4.8) [25]. Unilateral microinjections involved delivering 1.5 µl of dye-quenched fluorescein-conjugated gelatin (Life Technologies) intramolecularly at a rate of 0.25 µl/min. After the injections, there is a 30-minute incubation period during which the rats were either subjected to a sham or a stress procedure. MMP-2/9 gelatinase activity induces a proteolytic degradation of the gelatin, resulting in an activity-dependent increase in green fluorescence.

### CSF-1R inhibition via intraperitoneal injection of PLX3397

The majority of publications inhibiting microglia with PLX3397 (MedChemExpress, HY-16749, Monmouth Junction, NJ USA) are giving the drug via their normal chow for at least 1 week [26, 27]. Recent studies have used i.p. injections at a dose of 50 mg/kg to induce microglial ablation [28]. We chose not to ablate microglia, as doing so would create an artificial brain environment. Instead, animals received 1 mg/kg every 10-12 hours, mimicking consistent chow intake and following the treatment regimen previously used with i.p. injections [28]. The drug was diluted in 5% DMSO(Thermo Fisher, J66650.AK Waltham, MA, USA), 20% Kolliphor RH40 (Sigma Aldrich; C5135-500G, St. Louis, MI USA) in 1x phosphate buffer saline (PBS; Thermo Fisher, J62036.K3 Waltham, MA, USA). Animals were injected with either the drug or vehicle one week before the stress event, with the final injection given on the morning of the stress or sham session.

### Immunofluorescent staining

NAcore tissue slices and immunofluorescent staining were performed as described previously (Taborda-Bejarano et al). Briefly, microglia staining for morphology was performed by incubating an IBA-1 primary antibody (rabbit anti-IBA-1 1:300; Invitrogen PA5-21274, Waltham, MA, USA) for 20-24 hours at 4 °C, followed by a 2-hour room-temperature incubation with secondary antibody 568 anti-rabbit (1:200; Invitrogen, A-11011,Waltham, MA, USA). Microglia staining for CD68 colocalization used a different protocol. This protocol requires in-slide staining instead of free-floating staining. Once the tissue was adhered to the slide, it was photobleached using a blue light bulb. Tissue was then washed with 1X PBS for 5 minutes 3 times. For antigen retrieval, the tissue is submerged in 10mM sodium citrate and 0.05% tween 20 on MilliQ water for 20 minutes at 95 °C. The tissue is then washed and incubated in block solution (1x PBS with 0.5% Triton X-100 (0.5% PBST; Fisher Scientific AAA16046AP, Pittsburgh, PA USA) and 5% normal goat serum (Invitrogen, PI31873) for 1 hour at room temperature. Primary antibodies IBA-1(rabbit anti-IBA-1 1:300; Invitrogen PA5-21274, Waltham, MA, USA) and CD68(mouse anti-CD68 1:300; Abcam ab31630, Waltham, MA, USA) were diluted in block solution for 24 h at 4 °C. Followed by Secondary antibody incubation, Alexa 488 goat anti-rabbit and Alexa 633 goat anti-mouse (1:200, Thermofisher, A-21052, Waltham, MA, USA) in PBS.

### Confocal imaging

Microglia morphology within the NAcore was imaged using an unbiased protocol previously described in Taborda et al. (2025). Briefly, we use a Leica SP8 Upright Confocal Microscope (Mannheim, Germany) for our imaging. For each animal, we collect 6 images, 3 per stained brain hemisphere, at 63x magnification with a z-step size of 0.5, resulting in 76 z-steps. Brain regions with the zymography gel were imaged at 10x magnification, and ImageJ was used to quantify fluorescence.

### Data analysis

All data analyses were conducted using GraphPad Prism 11 (Boston, MA, USA). Prior to statistical testing, datasets were evaluated for normality using the D’Agostino & Pearson test. Non-normally distributed data, such as microglial morphology and microglial CD68 expression, were analyzed with non-parametric tests, specifically the Kruskal-Wallis test followed by Dunn’s post hoc test for multiple comparisons. qPCR data were processed using the delta-delta Ct method, with all data normalized to the Ct of the loading control and then further normalized to the difference in Ct between the gene of interest and the loading control in sham rats. These fold changes were analyzed using two-way ANOVA. Zymography data included multiple replicates per animal; for analysis, the six replicates with the highest fluorescence were selected and averaged per animal. These values were then subjected to two-way ANOVA. All analyses were performed under blind conditions.

## 1. Results

### CSF1R inhibition attenuates stress-induced MMP-2/9 activity in the NAcore

To determine whether microglia contribute to stress-induced MMP-2/9 activation, male and female rats were treated with the CSF1R inhibitor PLX3397 (PLX) prior to exposure to a single restraint stress session, and in vivo gelatinase activity, an indirect measure of MMP-2/9 activity, was quantified in the NAcore using fluorescent zymography (**Fig. 1A**). Animals were implanted with a unilateral NAcore-targeted cannula for gelatin microinjection and subsequent assessment of MMP-2/9 activity in one hemisphere (**Fig. 1**), while the contralateral hemisphere was collected for microglial inflammatory analyses (**Figs. 2**). Representative images of gelatinase activity of males rats are shown in **Figure 1B**.

**Figure 1.**
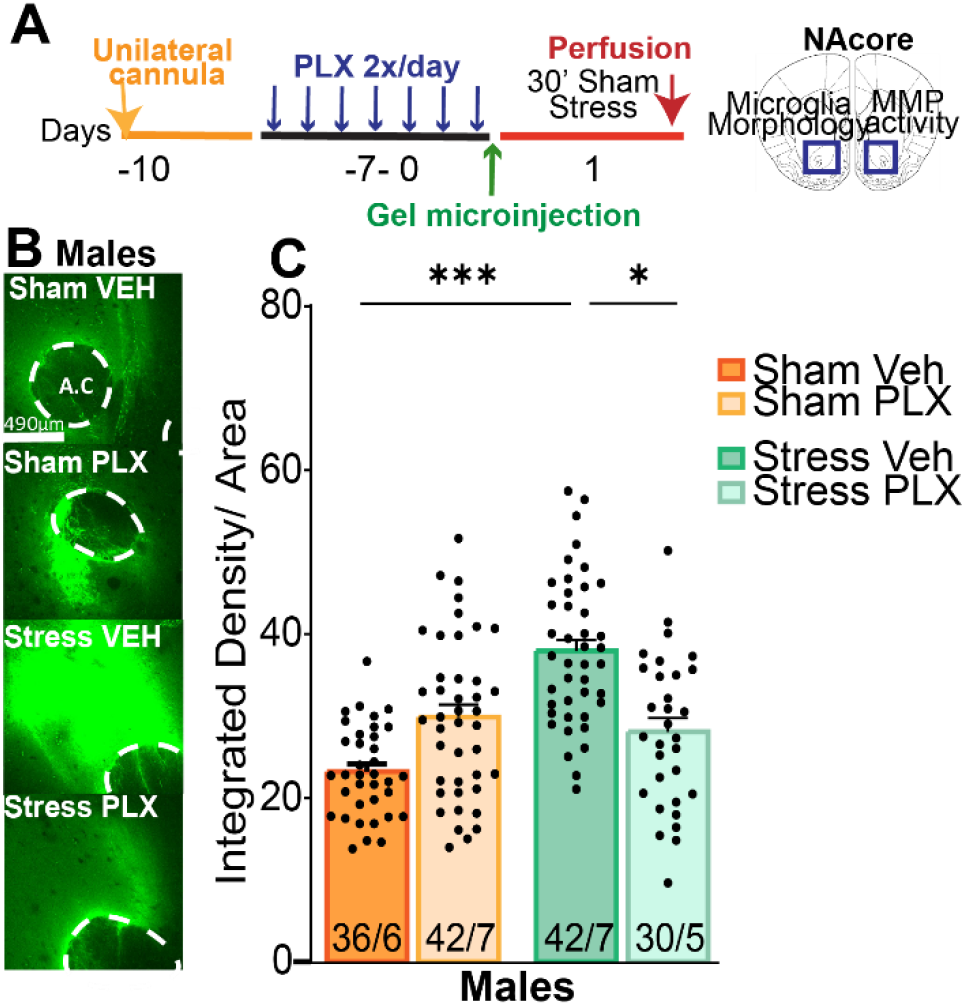
Stress-induced MMP-2/9 activity in the NAcore is attenuated by CSF1R inhibition. **A)** Experimental timeline illustrating unilateral cannula implantation into the NAcore, followed by twice-daily administration of the CSF1R inhibitor PLX3397 1mg/kg or vehicle (20% castor oil, 5% DMSO in PBS). Animals received in vivo microinjection of a fluorescent gelatin substrate, then brains were collected by perfusion immediately after the 30 minutes restraint stress experimental manipulation. **B)** Representative confocal images of in vivo zymography signal (green) in the NAcore of female and male, under sham and stress conditions in vehicle (VEH)- and PLX-treated animals. Dashed circles indicate the anterior commissure (A.C.). Scale bar = 490 μm. **C)** Quantification of MMP-2/9 activity in the NAcore across experimental groups. Acute stress significantly increased MMP-2/9 activity compared to sham controls, and this effect was attenuated by PLX treatment. Male: two-way ANOVA revealed an overall effect of stress and PLX (F(_1,21)_=11.51, *p*=0.003), with an increase in zymography fluorescence after stress (stress effect: F_(1,21)_=7.724, *p*=0.011, η^2^=18.44%, Sham VEH/Stress VEH p=0.0002), an effect that was blocked by PLX-induced CSF-1R inhibition (Stress VEH/ Stress PLX: p= 0.0118). Data are presented as individual data points overlaid on bar graphs (mean ± SEM). Numbers within bars represent number of images analyzed/number of animals).

**Figure 2.**
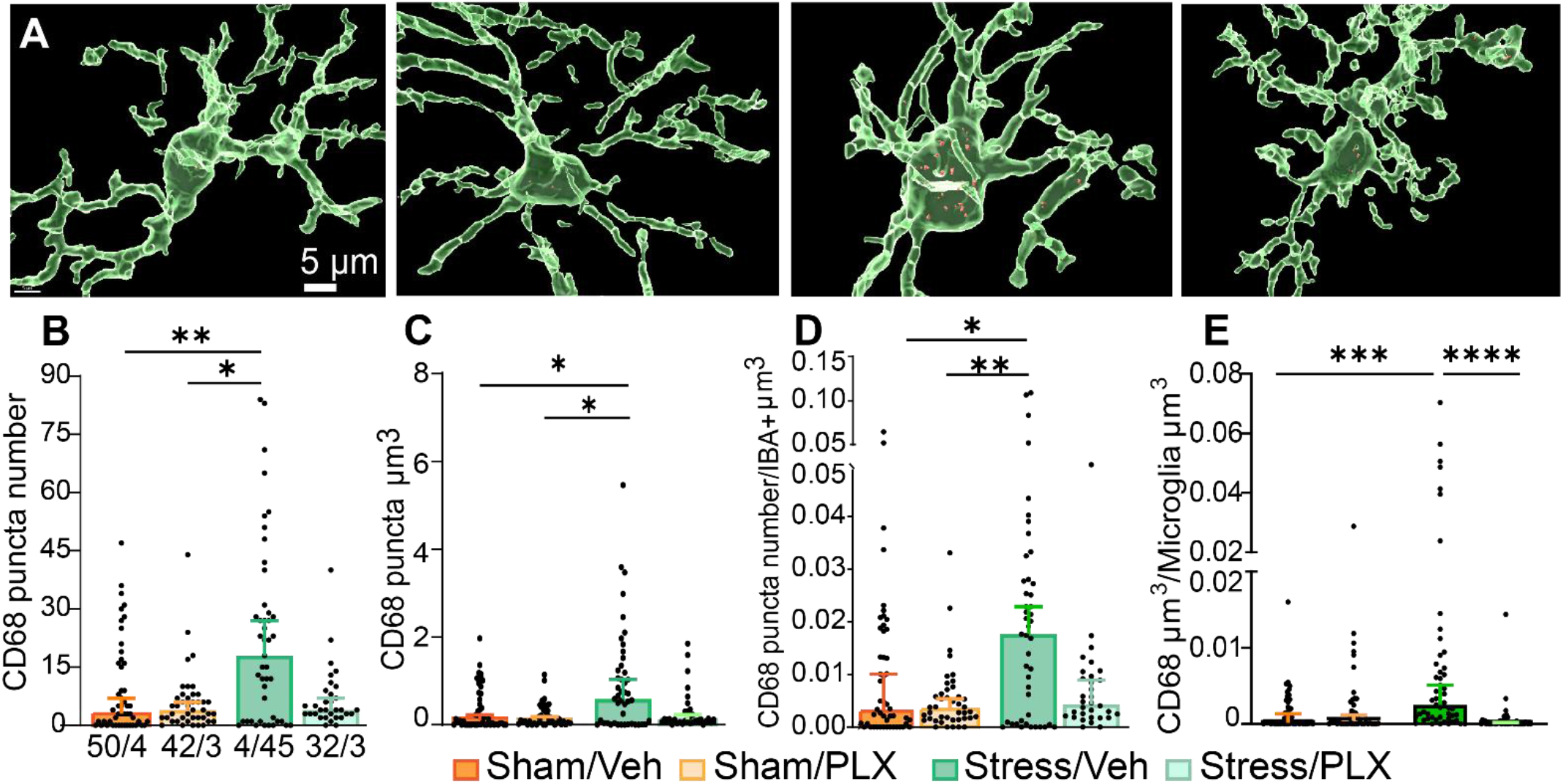
Stress-induced CD68 lysosomal phagocytic marker are attenuated by CSF1R inhibition in the NAcore. **A)** Representative three-dimensional reconstructions of IBA1-positive microglia (green) with CD68 puncta (red) in the NAcore from female rats across the 4 treatment groups, sham and stress animals treated with vehicle (VEH) or the CSF1R inhibitor PLX3397 (PLX). Scale bar = 5 μm. **B)** Stress-induced increased CD68 puncta number compared to sham/VEH and sham/PLX treatment. KW=13.30, *p*=0.0040. **C)** Stress-induced increased CD68 puncta volume compared to sham/VEH and sham/PLX treatment. KW=11.58, *p*=0.009. **D)** Stress-induced increased CD68 puncta number over IBA+ cells volume ratio compared to sham/VEH and sham/PLX treatment. KW=12.36, *p*=0.006. **E)** Stress-induced increased CD68 puncta volume over IBA+ cells volume ratio compared to sham/VEH and sham/PLX treatment. KW=12.36, *p*=0.006. Individual data points are overlaid on bar graphs representing median with 95% CI (data not normally distributed). Numbers within bars represent amount of cells/number of animals.

Acute stress increased MMP-2/9 activity in the NAcore of both male and female rats. PLX3397 (Pexidartinib) is a selective and potent small-molecule tyrosine kinase inhibitor that primarily targets the colony-stimulating factor 1 receptor (CSF1R) [26]. Inhibition of CSF1R signaling with PLX attenuated the stress-induced increase in MMP-2/9 activity in both males (**Fig. 1C**), indicating that microglial signaling contributes to stress-evoked MMP-2/9 activation and ECM remodeling within the NAcore.

### Stress increases microglial CD68, partially attenuated by CSF1R inhibition

To assess microglial recruitment following acute stress exposure, CD68 immunoreactivity was quantified in the NAcore 30 minutes after stress or sham exposure from the tissue processed from **Figure 1** (**Fig. 2A–F**). CD68 is a lysosomal-associated marker commonly used as an indicator of microglial phagocytic and inflammatory activity. Due to the non-normal distribution of microglial morphological data, non-parametric statistical analyses were performed. Representative three-dimensional reconstructions of IBA1-positive microglia with CD68-positive puncta from male are shown in **Figure 2A**.

Acute stress significantly increased microglial CD68 puncta number and volume in both male rats (**Fig. 2B, C**). To account for differences in microglial size, CD68 parameters were normalized to total microglial volume. This normalization yielded similar results, with stressed males showing significant increases in CD68 puncta number per microglial volume (**Fig. 2D**) and CD68 volume per microglial volume (**Fig 2E**). Together, these findings indicate that acute stress rapidly enhances microglial phagocytic associated signaling within the NAcore. Inhibition of CSF1R signaling with PLX attenuated the stress-induced increase in CD68. PLX treatment significantly reduced stress-induced CD68 parameters, in some cases decreasing values below sham control levels, suggesting robust suppression of stress-induced microglial activation.

### Acute stress induces a pro-inflammatory microglial-associated transcriptional profile in the NAcore

To further characterize microglial responses following acute stress exposure, we quantified microglial-associated and inflammatory-related mRNA expression in whole NAcore tissue using quantitative PCR (qPCR) (**Fig. 3**). While immunohistochemical analyses revealed rapid stress-induced changes in microglial CD68-associated activity, these approaches alone cannot fully capture the complexity of microglial signaling states. Therefore, we measured transcript levels associated with microglial signaling and inflammatory activation to better define the neuroimmune profile induced by 2 hours acute restraint stress.

**Figure 3.**
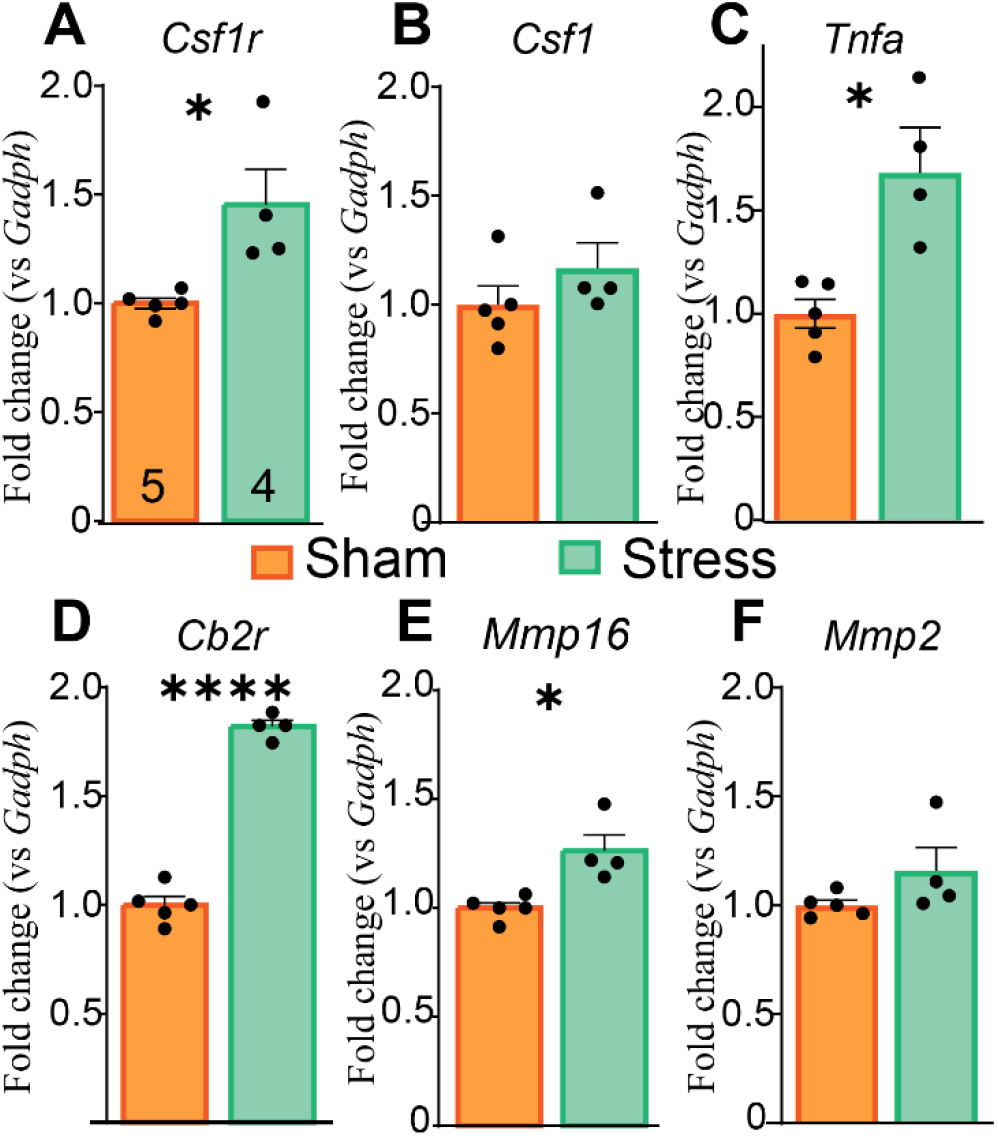
Stress-induced sex-dependent changes in microglial-associated inflammatory gene expression in the NAcore. Rats were stressed for 2 hours and then returned to their home cage for 1 hour, after which the animals were euthanized and dissected. **A-F)** Quantification of relative mRNA expression levels of microglial- and inflammatory-associated genes in the NAcore from male animals following sham or stress exposure. Gene expression was measured by qPCR and normalized to housekeeping genes, *Gadph*. **A)** *Csf1r* expression was increased following stress exposure (unpaired t test, t_7_=3.134 *p*=0.016). **B)** *Csf1* expression showed no significant differences between sham and stress groups. **C)** *Tnfa* expression was increased following stress exposure (unpaired t test, t_7_=3.264 *p*<0.0138. **D)** *Cb2r* expression was increased following stress exposure (unpaired t test, t_7_=16.27 *p*<0.0001). **E)** *Mmp16* expression was increased following stress exposure (unpaired t test, t_7_=2.43 *p*=0.435). **F)** *Mmp2* expression remained unchanged following stress exposure. Data are presented as individual data points overlaid on bar graphs representing mean ± SEM. Numbers within bars indicate n per group. Statistical comparisons were performed between sham and stress groups within each sex.

In male rats, acute stress significantly increased *Csf1r* mRNA expression (**Fig. 3A**), suggesting enhanced activation of CSF1R-dependent microglial signaling pathways involved in microglial survival and inflammatory activation. In contrast, *Csf1*, one of the primary ligands for CSF1R, was not significantly altered following stress exposure (**Fig. 3B**). Acute stress also increased *Tnfa* expression (**Fig. 3C**), indicating enhanced pro-inflammatory cytokine signaling within the NAcore. Similarly, *Cnr2* (*Cb2r*), a cannabinoid receptor associated with immune regulation and microglial activation, was elevated following stress (**Fig. 3D**). In addition, stress significantly increased *Mmp16* expression **(Fig. 3E**), a membrane-associated matrix metalloproteinase involved in ECM remodeling and activation of other MMPs, while *Mmp2* transcript levels remained unchanged between sham and stress groups (**Fig. 3F**). These findings indicate that acute stress promotes a pro-inflammatory and ECM-associated transcriptional profile in the male NAcore.

Together, these findings indicate that acute restraint stress induces a strong microglial-associated inflammatory transcriptional response in the male NAcore, supporting stress-induced neuroimmune signaling.

## Discussion

### Microglial CSF1R signaling regulates stress-induced ECM remodeling

The present study demonstrates that acute restraint stress rapidly induces microglial-associated neuroimmune signaling and ECM remodeling within the nucleus accumbens core (NAcore), and that these adaptations are partially dependent on microglial CSF1R signaling. Specifically, we show that acute stress increases MMP-2/9 activity, enhances microglial CD68-associated phagocytic signaling, and promotes expression of genes associated with microglial activation, inflammatory signaling, and ECM remodeling. Moreover, pharmacological inhibition of CSF1R signaling with PLX3397 (1 mg/kg) attenuated stress-induced MMP-2/9 activation and reduced stress-induced increases in CD68-associated signaling, supporting a role for microglia as upstream regulators of stress-induced ECM remodeling.

These findings extend our previous work demonstrating that stress and psychostimulants engage common mechanisms of synaptic plasticity within the NAcore. MMP-2 and MMP-9 are well-established regulators of ECM remodeling and dendritic spine plasticity[9, 10, 29-32]. Previous studies from our laboratory demonstrated that MMP-2/9 activity is required for cocaine-induced synaptic adaptations and reward-associated behaviors[9, 10]. The present findings indicate that acute stress similarly recruits MMP-dependent remodeling mechanisms and identify microglia as a critical upstream component of this process. Because inhibition of CSF1R signaling attenuated stress-induced MMP-2/9 activity, these results suggest that active microglial signaling is required for stress-induced ECM remodeling within reward circuitry.

Emerging evidence identifies microglia as central regulators of ECM homeostasis and remodeling[14, 33]. Microglia contribute to ECM turnover through multiple complementary mechanisms, including the secretion of matrix metalloproteinases and the phagocytic clearance of extracellular substrates[33]. In particular, microglia are a recognized source of MMP-2 and MMP-9, enzymes capable of degrading ECM proteins that regulate synaptic stability and plasticity[34]. The ability of acute stress to increase MMP-2/9 activity, coupled with the attenuation of this response by CSF1R inhibition, suggests that stress recruits a microglial remodeling program that promotes extracellular proteolysis within the NAcore[33]. Consistent with this interpretation, previous studies have demonstrated that CSF1R inhibition or microglial depletion reduces ECM degradation and preserves ECM structures, supporting a fundamental role for microglia in regulating matrix integrity[35, 36]. Together, these findings position microglial CSF1R signaling as a critical upstream regulator of stress-induced ECM remodeling.

### Acute stress enhances microglial phagocytic-associated signaling

Acute stress robustly increased CD68-associated signaling within NAcore microglia. CD68 is a lysosomal-associated protein commonly used as a marker of phagocytic activity and cellular engagement[37]. Increased CD68 puncta number, volume and ratio indicate that microglia rapidly transition to an activated functional state following stress exposure. Importantly, these effects were attenuated by CSF1R inhibition, demonstrating that stress-induced microglial engagement depends, at least in part, on CSF1R signaling. Collectively with the zymography findings, these results support a model in which stress recruits microglial signaling pathways that contribute to ECM remodeling.

Beyond its role as a marker of microglial activation, accumulating evidence indicates that CD68-positive lysosomal compartments participate directly in ECM engulfment and degradation. Microglia are increasingly recognized as active regulators of ECM remodeling through two complementary mechanisms: proteolytic degradation mediated by enzymes such as MMP-2 and MMP-9 and phagocytic clearance of ECM components[33]. Thus, the concurrent increases in CD68-associated signaling and MMP-2/9 activity observed following stress suggest that microglia rapidly adopt a remodeling-associated phenotype characterized by enhanced lysosomal activity, phagocytosis, and extracellular proteolysis. The observation that CSF1R inhibition attenuated both CD68-associated signaling and MMP-2/9 activation further supports a coordinated microglial remodeling response to stress.

This interpretation extends beyond a purely inflammatory view of microglial function. Rather than acting solely as immune effector cells, microglia are now recognized as key regulators of synaptic plasticity through their ability to modify the ECM and shape the perisynaptic environment. By regulating ECM composition through both protease release and phagocytic remodeling, microglia can influence dendritic spine dynamics, synaptic stabilization, and neuronal communication. Consequently, the stress-induced increases in CD68 and MMP-2/9 activity observed in the present study likely represent early cellular events that initiate ECM remodeling and create a permissive environment for subsequent synaptic plasticity within the NAcore.

### Acute stress induces a microglial-associated inflammatory and ECM transcriptional profile

To further characterize stress-induced neuroimmune signaling, we quantified expression of genes associated with microglial activation, inflammatory signaling, and ECM remodeling. Acute stress increased expression of Csf1r, suggesting enhanced activation of CSF1R-dependent signaling pathways involved in microglial responsiveness and inflammatory regulation, similar to what it has been shown following chronic stress [38]. Moreover, pharmacological blockade of glucocorticoid receptor suppresses chronic stress-induced CSF1 signaling in the prefrontal cortex, reduced microglial phagocytic markers, and prevented stress-related cognitive impairments [39]. CSF1R/CSF1 signaling is amplified under inflammatory conditions [39,40], underscoring its sensitivity to stress-related immune activation. To define the cellular sources of CSF1R signaling relevant to microglia–neuron crosstalk, we analyzed mRNA expression using publicly available single-nucleus RNA-sequencing data from the adult rat NAc under baseline conditions (saline treated rats)[40] also see development [41]. RATLAS website[40] search indicate that *Csf1r* expression was highly enriched in microglia compared with astrocytes (adjusted *p*=9.39×10−^116^ vs. *p*=0.035) and any other cell population, confirming that CSF1R signaling capacity is largely restricted to microglia. In contrast, expression of Csf1 remained unchanged, indicating that stress-induced activation of the CSF1R pathway may occur through increased receptor expression, altered receptor sensitivity, or alternative ligands such as IL-34[21, 22]..

Stress also increased expression of *Tnfa*, consistent with enhanced pro-inflammatory cytokine signaling within the NAcore. TNFα is a potent regulator of microglial-neuronal communication and has been implicated in synaptic remodeling[15, 16], but also see [17] and MMP regulation[18, 19]. In addition, stress increased expression of Cnr2, the gene encoding the cannabinoid receptor CB2, a receptor highly associated with immune regulation and activated microglial states[42]. Elevated *Cnr2* expression may reflect compensatory immune-regulatory mechanisms recruited in response to stress-induced neuroimmune activation.

Finally, stress increased expression of Mmp16, a membrane-associated metalloproteinase that contributes to ECM turnover and activation of downstream MMP signaling pathways. In contrast, Mmp2 transcript levels were unchanged despite robust increases in MMP-2/9 enzymatic activity. This dissociation suggests that stress-induced gelatinase activity is likely regulated through post-transcriptional mechanisms, activation of existing enzyme pools, or upstream signaling pathways rather than increased Mmp2 transcription. Together, these findings support the conclusion that acute stress promotes a coordinated neuroimmune and ECM remodeling response within the NAcore.

### Study limitations

Several limitations should be considered. First, qPCR analyses were performed using whole NAcore tissue punches and therefore do not provide cell-type-specific resolution. Consequently, the observed transcriptional changes cannot be definitively attributed to microglia. Future studies using single-cell RNA sequencing, fluorescence-activated cell sorting, or spatial transcriptomic approaches will be necessary to identify the cellular sources of these signals.

Second, although PLX3397 is widely used to target CSF1R signaling, it may also affect other tyrosine kinases, including c-Kit and FLT3. Thus, some effects cannot be exclusively attributed to CSF1R inhibition. Future studies using microglia-specific genetic approaches will help establish causal cellular mechanisms.

Third, the present study focused on acute stress responses at a single post-stress time point. Because microglial signaling and ECM remodeling are highly dynamic processes, future work should examine the temporal progression of these adaptations and determine how they contribute to long-lasting synaptic plasticity and behavioral outcomes.

Finally, although the present findings strongly support a role for microglia in stress-induced ECM remodeling, the precise molecular mechanisms linking CSF1R activation to MMP-2/9 signaling remain to be established. Future studies will determine whether cytokine-dependent signaling pathways, including TNFα-mediated mechanisms, directly couple microglial activation to ECM remodeling and synaptic plasticity.

## Conclusion

In summary, acute restraint stress rapidly increases MMP-2/9 activity, microglial CD68-associated signaling, and expression of genes associated with microglial activation, inflammatory signaling, and ECM remodeling within the NAcore. Pharmacological inhibition of CSF1R signaling attenuated both stress-induced MMP activation and microglial engagement, identifying microglia as critical upstream regulators of ECM remodeling. Together with emerging evidence that microglia coordinate matrix turnover through both proteolytic and phagocytic mechanisms, our findings support a model in which stress recruits a CSF1R-dependent microglial remodeling phenotype that initiates ECM reorganization within reward circuitry. These data provide mechanistic insight into how acute stress engages neuroimmune pathways to shape synaptic plasticity and may contribute to maladaptive circuit adaptations associated with neuropsychiatric disorders.

